# VEGF-C overexpression in kidney progenitor cells is a model of renal lymphangiectasia

**DOI:** 10.1101/2023.05.03.538868

**Authors:** Michael D. Donnan, Dilip K. Deb, Valentin David, Susan E. Quaggin

**Author notes:** **Correspondence:** Michael Donnan Tele.

## Abstract

**Background:** Lymphangiogenesis is believed to be a protective response in the setting of multiple forms of kidney injury and mitigates the progression of interstitial fibrosis. To augment this protective response, promoting kidney lymphangiogenesis is being investigated as a potential treatment to slow the progression of kidney disease.

As injury related lymphangiogenesis is driven by signaling from the receptor VEGFR-3 in response to the cognate growth factor VEGF-C released by tubular epithelial cells, this signaling pathway is a candidate for future kidney therapeutics. However, the consequences to kidney development and function to targeting this signaling pathway remains poorly defined.

**Methods:** We generated a new mouse model expressing *Vegf-C* under regulation of the nephron progenitor Six2Cre driver strain (*Six2Vegf-C*). Mice underwent a detailed phenotypic evaluation. Whole kidneys were processed for histology and micro computed tomography 3-dimensional imaging.

**Results:** *Six2Vegf-C* mice had reduced body weight and kidney function compared to littermate controls. *Six2Vegf-C* kidneys demonstrated large peripelvic fluid filled lesions with distortion of the pelvicalcyceal system which progressed in severity with age. 3D imaging showed a 3-fold increase in total cortical vascular density. Histology confirmed a substantial increase in LYVE1+/PDPN+/VEGFR3+ lymphatic capillaries extending alongside EMCN+ peritubular capillaries. There was no change in EMCN+ peritubular capillary density.

**Conclusions:** Kidney lymphangiogenesis was robustly induced in the *Six2Vegf-C* mice. There were no changes in peritubular blood capillary density despite these endothelial cells also expressing VEGFR-3. The model resulted in a severe cystic kidney phenotype that resembled a human condition termed renal lymphangiectasia. This study defines the vascular consequences of augmenting VEGF-C signaling during kidney development and provides new insight into a mimicker of human cystic kidney disease.

## INTRODUCTION

Chronic kidney disease is accompanied by progressive changes in the kidney microvasculature which subsequently influence disease progression. The peritubular capillaries undergo injury-related rarefaction and dropout which can contribute to tissue hypoxia and interstitial fibrosis.^1^ Conversely, kidney lymphatic capillaries undergo expansion, termed lymphangiogenesis, in the setting of multiple forms of kidney disease including IgA nephropathy,^2^ diabetic kidney disease,^3^ renal fibrosis,^4^ renal cystogenesis,^5^ and transplant rejection^6^. Recent evidence has suggested lymphangiogenesis is a protective response and attenuates the progression of fibrosis.^7^ Additionally, augmented lymphangiogenesis was demonstrated to be protective in early animal models of acute kidney injury, and as such the therapeutic promotion of lymphangiogenesis has been proposed as a new avenue for the treatment of kidney disease.^8,9^

The primary driver of lymphangiogenesis is the activation of the tyrosine kinase receptor VEGFR-3 (also known as FLT4) expressed on the surface of LECs.^10-12^ The cognate growth factors VEGF-C, and to a lesser extent VEGF-D, are the primary ligands for VEGFR-3 signaling.^11,13,14^ These growth factors are released by circulating macrophages and kidney tubular epithelial cells in response to injury.^15-17^ VEGF-C induced lymphangiogenesis is observed across multiple modalities of animal kidney injury models.^16-18^ Supporting the potential therapeutic benefit of augmenting lymphangiogenesis in kidney injury, either transgenic overexpression or exogenous administration of VEGF-C or -D was protective in rodent models of acute kidney injury^4,8^, transplant rejection^19^, and polycystic kidney disease.^5^

However, we have recently shown that VEGFR-3 expression is not exclusive to lymphatic endothelial cells in the kidney and that targeting this signaling pathway may have unexpected off-target consequences. VEGFR-3 is also expressed by the fenestrated blood capillaries of the kidney where it regulates glomerular capillary development.^20^ This role of VEGFR-3 was limited to a critical mid-embryonic developmental period and was independent of paracrine VEGF-C signaling highlighting the time-and-cell dependent heterogeneity of signaling between endothelial populations. While targeting VEGF-C / VEGFR-3 signaling appears to be promising for the development of new therapeutics to treat kidney diseases it remains critical to first determine the intended and off-target consequences of augmenting VEGF-C expression within the kidney to guide rational therapeutic design.

In this study, we used a VEGF-C gain-of-function model (*VegfcGOF*) to determine the effect of increased *Vegf-C* expression on kidney lymphangiogenesis and to evaluate for off-target consequences to kidney blood vascular morphology and kidney function. Here we show that VEGF-C has heterogeneous effects on the VEGFR3+ endothelial populations within the kidney and that overexpression during development results in a severe cystic phenotype reminiscent of renal lymphangiectasia.

## RESULTS AND DISCUSSION

To study the effect of augmenting VEGF-C signaling within the kidney we generated a mouse model (*Six2Vegf-C*) using the *VegfcGOF* and *Six2cre* mouse lines (Figure 1A). In this model, *Vegfc* is overexpressed in the SIX2+ kidney nephron progenitor cells as a method to mimic the tubular epithelial cell expression pattern of *Vegfc* observed during injury related lymphangiogenesis.^16^ *Six2Vegf-C* mice were runted in comparison to Vegfc^EF1-/-^ : Six2Cre^+/-^ littermate controls (Ctrl mean weight 26.2 g, Six2Vegf-C mean weight 14.57 g, difference -11.63 g, SEM ± 1.111 g, p=0.001) and had reduced viability typically limited to 4-8 weeks of age (Figure 1B-C). Strikingly, these mice had enlarged, fluid filled kidneys of variable severity that were consistently associated with a reduction in kidney function (Ctrl mean BUN 25.73 mg/dL, Six2Vegf-C mean BUN 71.22 mg/dL, difference +45.50 mg/dL, SEM ± 5.58 md/dL, p<0.0001). No other phenotype was observed in gross histology suggesting the runted size and limited viability of these mice was due to renal insufficiency.

**Figure 1:**
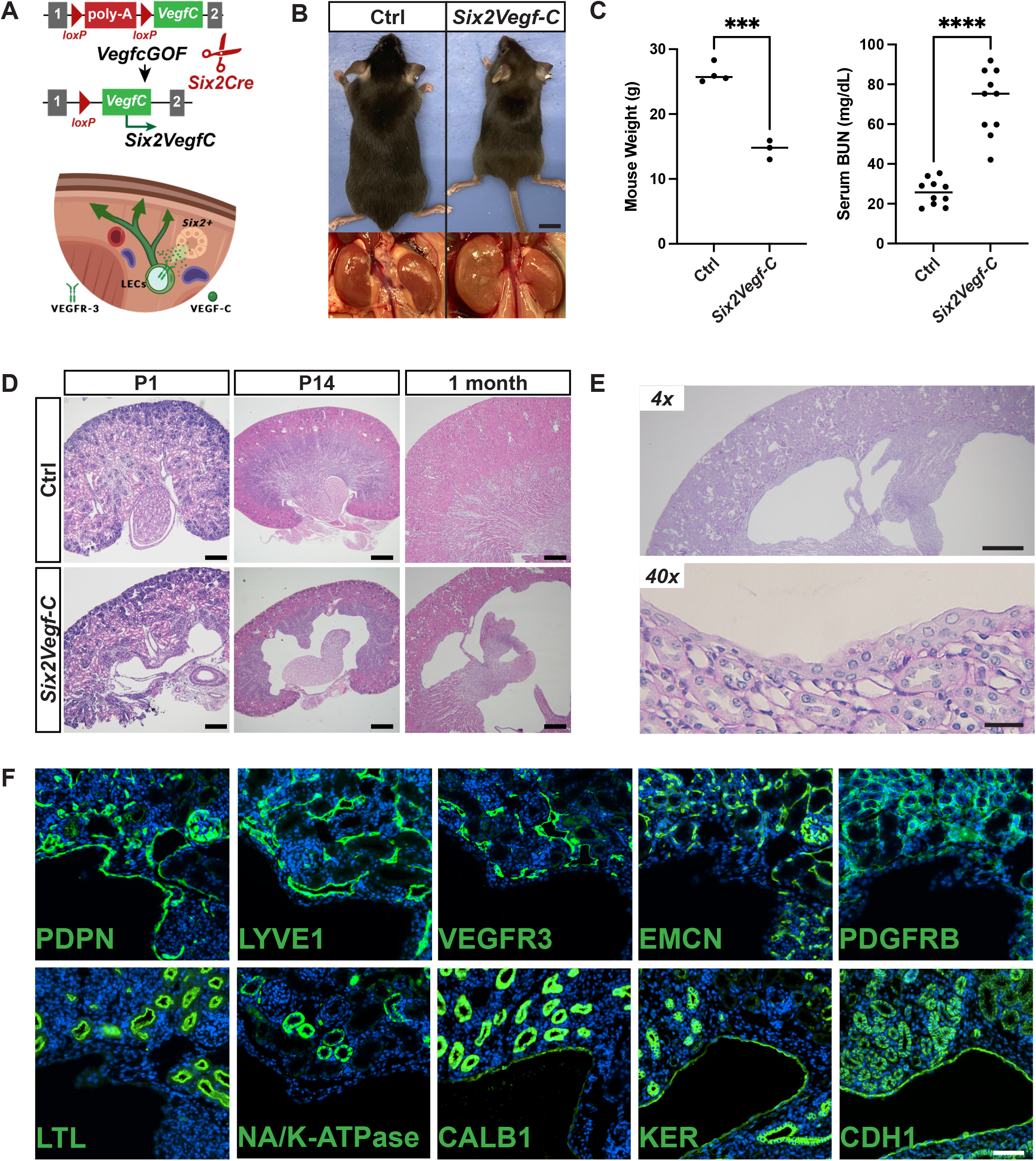
*Six2Vegf-C* mice develop large cystic malformations resembling renal lymphangiectasia. **A**, Schematic representation of the *Six2VegfC* mouse model where *Six2*+ nephron progenitor cells overexpress the growth factor VEGF-C which interacts with lymphatic endothelial cells (LECs) to promote lymphangiogenesis during kidney development. **B**, Representative images of *Six2Vegf-C* mice and littermate controls (Ctrl). *Six2Vegf-C* mice appear runted and have enlarged appearing kidneys of varying severity. Scale bar: 1 mm. **C**, Mouse weight in grams (g) and serum blood urea nitrogen (BUN) demonstrate reduced weight and kidney function in *Six2Vegf-C* mice. All data are presented as ±SE. Statistical comparisons were made using an unpaired t test. ***P = 0.0001 and ****P < 0.0001. **D**, Representative images of H&E stained kidney sections from *Six2Vegf-C* mice and littermate controls at postnatal day 1 (P1), P14, and 1 month of age demonstrating progressive peripelvic cystic lesions. Scale bar: 200 μm (P1), 500 μm (P14), 500 μm (1 month). **E**, Higher magnificent images of PAS stained kidney sections from *Six2Vegf-C* mice. Scale bar: 500 μm (4x), 25 μm (40x). **F**, Immunofluorescence of *Six2Vegf-C* kidney cystic lesions labeled for podoplanin (PDPN), LYVE1, VEGFR3, endomucin (EMCN), PDGFRB, Lotus tetraglobulus lectin (LTL), NA/K-ATPase, calbindin 1 (CALB1), pan-keratin (KER), cadherin-1 (CHD1). Scale bar: 50 μm. N ≥ 3 animals/group for all representative images.

On H&E histology, *Six2Vegf-C* mice demonstrate dramatic cyst-like lesions within the renal pelvis. These lesions are observed by post-natal day 1 (P1) and grow in severity with kidney maturity (Figure 1D). By 1 month of age, there is marked distortion of the pelvicalyceal system with loss of corticomedullary differentiation. These lesions correlate with the enlarged fluid filled appearance of *Six2Vegf-C* mouse kidneys on gross histology and resemble human renal lymphatic malformations termed renal lymphangiectasia. Renal lymphangiectasia also present as peripelvic fluid filled lesions often distorting the pelvicalyceal system and can be often misidentified for other cystic kidney lesions including hydronephrosis, cystic nephromas, and occasionally polycystic kidney disease.^21^ The exact cause of this condition is not known but both familial and acquired forms exist; the latter can be seen in patients with renal vein thrombosis or after kidney transplantation.^22,23^ It is suspected that either functional or mechanical disruption to the typical lymphatic drainage through the large lymphatic trunks within the renal hilum precipitates lymphangiectasia.

Higher magnification of Periodic Acid–Schiff-stained images of the outer wall of these cystic lesions demonstrates an irregular layer of flattened and cuboidal cells (Figure 1E). To determine the origin of these lesions, kidney sections were processed for immunofluorescent imaging (Figure 1F) with markers for lymphatic endothelial cells (PDPN, LYVE1, VEGFR3), blood endothelial cells (VEGFR3, EMCN), mesenchymal cells (PDGFR-β), and epithelial cells (LTL, NA/K-ATPase, CALB1, KER, CDH1). While the inner lining of these cystic lesions had strong expression of podoplanin (PDPN), they lacked co-expression with other expected lymphatic endothelial cell markers (LYVE1 and VEGFR3). Rather, the cyst wall expressed calbindin-1 (CALB1), pan-keratin (KER), and cadherin-1 (CDH1) consistent with the epithelial cells of the kidney distal tubules and collecting ducts. Together, this suggests these lesions are less likely to be primarily large lymphatic malformations but a dilation of the kidney collecting system more consistent with hydronephrosis.

Clinically, it can be difficult to distinguish between hydronephrosis and suspected renal lymphangiectasia.^24^ Both conditions present as large peripelvic fluid collections while renal lymphangiectasia can be distinguished by contrast enhanced Computed Tomography (CT) imaging lacking opacification of the cystic fluid separating the lesion from the urinary calyceal system. However, there can be an overlap between the two pathologies. Renal lymphatic malformations are reported to lead to hydronephrosis through compression and/or disruption of the calyceal system which passes in parallel to the lymphatic system through the kidney hilum.^21^ In the developing mouse kidney, the lymphatic system originates from a plexus located in the kidney hilum.^25,26^ In contrast to injury associated lymphangiogenesis in the adult kidney, during development, *Vegf-C* is not expressed by kidney tubular epithelial cells.^20^ Instead, developmental *Vegf-C* expression is localized to the kidney arteriolar endothelial cells and likely directs the normal expansion of the hilar lymphatic plexus into the developing lymphatic vessels which run in parallel to the major vascular bundles.^20,27^ It is suspected that disruption of the typical gradient of VEGF-C from kidney arteriolar cells may adversely affect the organized expansion of the kidney hilar lymphatic plexus leading to compression of the calyceal system leading to the observed cystic phenotype.

Despite the atypical pattern of *Vegf-C* expression during development, *Six2Vegf-C* mice still develop lymphatic capillaries within the kidney cortex with a substantial increase in vascular density compared to littermate controls (Figure 2). In the P1 mouse kidney, lymphatic vessels are typically sparse in the kidney cortex. In contrast, *Six2Vegf-C* mice have extensive expansion of LYVE+ lymphatic capillaries throughout the kidney cortex primarily adjacent to the EMCN+ peritubular capillaries (Figure 2A). These capillaries express traditional lymphatic endothelial cell markers including LYVE1, PDPN, VEGF3, and NRP2 (Figure 2B). Micro-CT imaging of the kidney from 1 month old mice demonstrates the large peripelvic fluid collections with an increased density of disorganized vascular structure (Figure 2C). Cortical vascular density as calculated by micro-CT imaging is increased in *Six2Vegf-C* mice (Mean vessel number (VM) per mm^3^: *Six2Vegf-C* 21.16 VN/mm^3^, Ctrl 6.521 VN/mm^3^, mean difference 14.64 VN/mm^3^, SEM ± 1.253 VN/mm^3^, P = 0.0074) with a shift in distribution to larger diameter vessels compared to controls. The shift towards higher density of larger diameter vessels suggests the increase in vascular density was due to an expansion of lymphatic vessels (Figure 2D).

**Figure 2:**
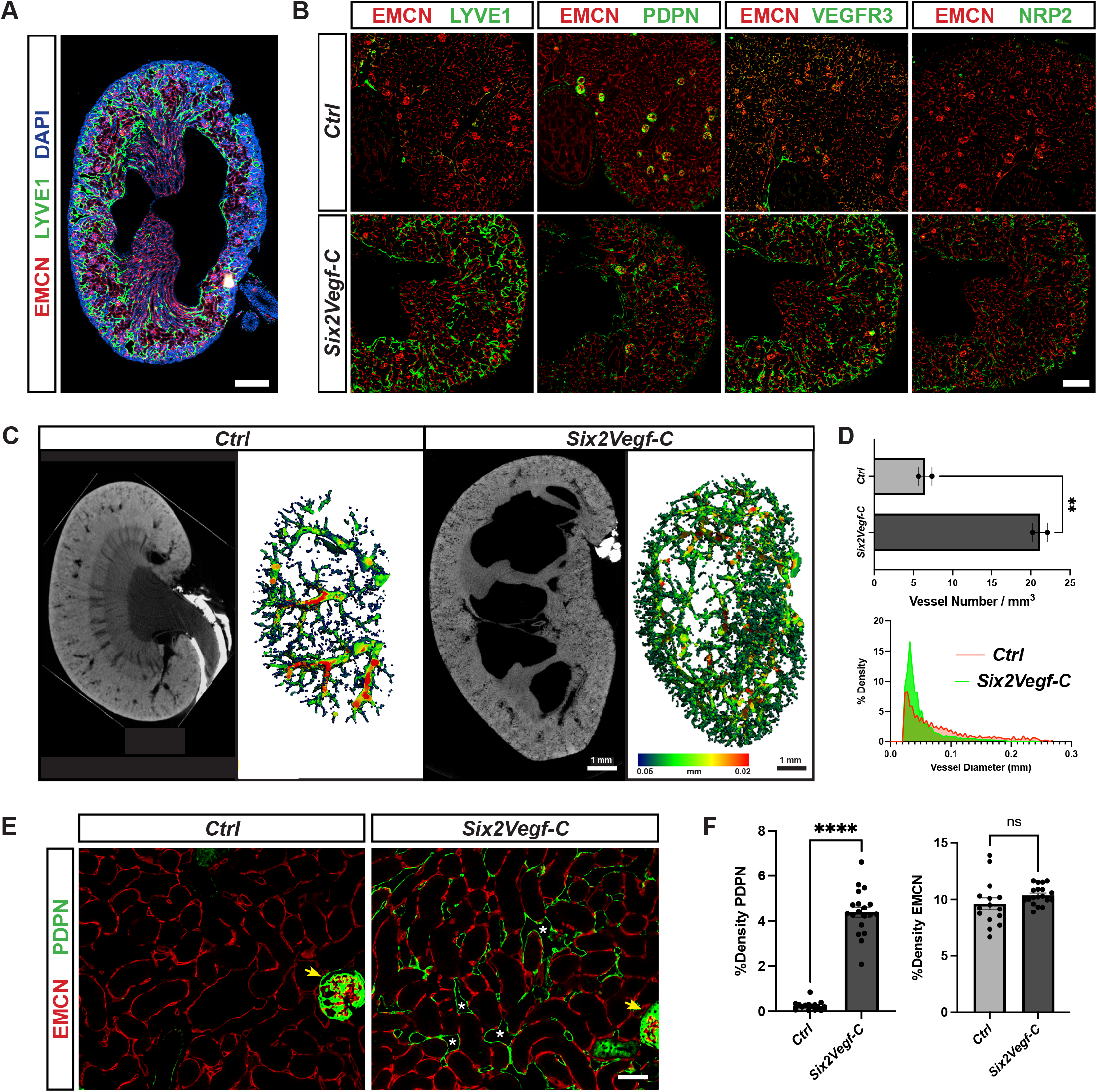
VEGF-C increases the density of lymphatic vessels in the kidney cortex without altering the peritubular capillaries. **A**, Representative image of a *Six2Vegf-C* P1 mouse kidney with a peripelvic cystic lesion and labeled for blood capillaries with endomucin (EMCN; red), lymphatic vessels with LYVE1 (green), and DAPI (blue). Scale Bar: 250 μm. **B**, *Six2Vegf-C* mice have an increased number of lymphatic vessels in the kidney cortex as compared to controls labeled by the LEC markers LYVE1, podoplanin (PDPN), VEGFR, and neuropilin 2 (NRP2) (Green). Sections are co-labeled with endomucin (EMCN; red). Scale Bar: 250 μm. n ≥ 3 animals/group for representative images. C, Contrast enhanced 3D microtomography images of whole mouse kidneys with vascular structure and heat-map representation of vascular diameter. Heat-map scale range from 0.05 mm (blue) through 0.2 mm (red). Scale bar: 1 mm. **D, Above:** Cortical vascular density in vessel number per mm^3^ as determined by analysis of microtomography images. Data are presented as ±SE. Statistical comparisons were made using an unpaired t test. **P = 0.0.0074. n = 2 animals/group. **Below**: Comparison of % Density against vessel diameter (mm) in the kidney cortex between *Six2Vegf-C* and control animals. **E**, Immunofluorescence of the mouse kidney cortex at 1 month of age as labled by endomucin (EMCN; red) and podoplanin (PDPN, green). White asterisk denotes large diameter lumen of PDPN+ lymphatic capillaries. Yellow arrow denotes extra-lymphatic labeling of glomerular podocytes by PDPN, seen similarly in both *Six2VEGF-C* and control groups. Scale Bar: 50 μm. n ≥ 6 animals/group for representative images. F, Kidney cortex vascular density as percentage of vasculature area to total area for lymphatic vessels (PDPN, left), and peritubular capillaries (EMCN, right). All data are presented as ±SE. Statistical comparisons were made using an unpaired t test. ****P < 0.0001 and ns = non-significant. n = 6 animals per group with 2-3 sections of kidney cortex evaluated per animal.

To further quantify the extent of lymphangiogenesis in *Six2Vegf-C* mice, we evaluated the change in PDPN+ vascular density in the kidney cortex. In the P14 mouse kidney cortex, lymphatic density is sparse with the marker PDPN primarily being observed only in the podocytes of the glomerulus (Figure 2E). *Six2Vegf-C* mice have a substantial increase in PDPN+ density (Mean percentage of PDPN+ cortical area: Ctrl 0.245%, Six2Vegf-C 4.394%, mean difference +4.148%, SEM ±0.26%, P <0.0001) (Figure 2F). Additionally, the diameter of intrarenal PDPN+ vessels appear enlarged, suggesting irregular or dilated lymphatic vessels. Dilated intrarenal lymphatic vessels are also observed in the setting of renal lymphangiectasia and may reflect impairment of lymphatic drainage from the kidney.

As VEGFR3 is also expressed in the fenestrated blood endothelial cells (BECs) of the kidney^20^, and VEGF-C can signal through VEGFR2 expressed by BECs^28^ we evaluated for changes in kidney peritubular capillary density (Figure 2E&F). The density of EMCN+ peritubular capillaries in the kidney cortex was not significantly different between *Six2Vegf-C* mice and controls (Mean percentage of EMCN+ cortical area: Ctrl 9.62%, Six2Vegf-C 10.37%, mean difference +0.755%, SEM ±0.5097%, P = 0.1483). Additionally, the morphology of EMCN+ peritubular capillaries was unchanged between groups. This supports that while there is an overlap between vascular growth factor signaling between blood and lymphatic endothelial cells, VEGF-C in the kidney primarily drives lymphangiogenesis without significantly altering the blood capillary density or morphology.

Together, this study highlights that kidney tubular epithelial cell expression of *Vegf-C* robustly promotes lymphangiogenesis in the mouse kidney without altering VEGFR3+ blood capillary density. This raises the possibility of targeting this pathway with future therapeutics to promote lymphangiogenesis for the treatment of kidney disease. However, altering kidney VEGF-C expression can lead to off-target consequences to kidney function as seen by the development of a severe cystic phenotype associated with renal insufficiency. As the pathophysiology of renal lymphangiectasias remains poorly defined, this model also provides new insight into the development of this rare condition.

## METHODS

### Mouse strains and husbandry

The *Vegfc*GOF mouse line has been previously described^29^ and was provided by the Oliver lab at Northwestern University (Chicago, Illinois). Briefly, the first intron of the *Eif1a* locus contains the full-length cDNA of mouse *Vegf-C*. A floxed triple poly(A) cassette precedes the *Vegf-c* cDNA preventing expression until removed via Cre-mediated excision. The Six2Cre mouse line was a gift from Dr. Andrew McMahon (University of Southern California, Los Angeles, CA) and has been previously described^30^. *VegfcGOF* mice were crossed with *Six2Cre* mice to create the *Six2Vegf-C* mouse line and induce kidney nephron progenitor expression of *Vegf-C*. Both male and female mice were used in all experiments and all data includes a combination of both sexes. Animals were genotyped by genomic PCR analysis using the following primers: EF1, forward 5′- CAGAAGACCGTGTGCGAATC-3′, reverse 5′- CGATTACGACGATGTTGATGT-3′; Cre, forward 5′- GTGCAAGTTGAATAACCGGAAATGG-3′, reverse 5′- AGAGTCATCCTTAGCGCCGTAAATCAAT-3′;. Mice were reared, bred, and characterized according to strict ethical guidelines approved by the Institutional Animal Care and Use Committee of Northwestern University.

### Blood Urea Nitrogen (BUN) assay

BUN assay was performed by the UAB-USCD O’Brien Center for Acute Kidney Injury Research. A quantitative colorimetric assay based on an improved Jung method from Bioassays Systems was used to measure urea. The chromogenic reagent formed a colored complex with urea and the intensity of the color, measured at 520 nm, was directly proportional to the urea concentration in the sample. Each sample is assayed in duplicate. BUN (mg/dL) = [Urea] / 2.14.

### Histology and Histochemistry

Tissues and organs were routinely fixed in 4% formaldehyde in phosphate buffered saline (PBS, pH7.5) for 24 hours at 4°C. Fixed tissues were embedded in paraffin blocks to produce 4- μm thick sections for routine histology (hematoxylin-eosin and periodic acid-aldehyde Schiff staining), and immunostainings. Standard methods for immunofluorescence processing were carried out following heat induced antigen retrieval by citrate buffer (0.01M; pH 6.0) using the following antibodies: PDPN (DSHB Cat# 8.1.1), LYVE1 (R&D Cat# AF2125), VEGFR3 (R&D Cat# AF743), NRP2 (R&D Cat# AF567 EMCN (Abcam Cat# ab106100), PDGFRB (R&D Cat# AF1042), LTL (*Lotus tetraglobulus lectin*) (Vector Laboratories Cat# FL-1321), NA/K-ATPase (DSHB Cat# a5), CALB1 (CST Cat# 13176S), pan-Keratin (CST Cat# 4545S), and CDH1 (CST Cat# 3195S). Fluorochrome-conjugated secondary antibodies were used appropriately. For CALB1, CDH1, and pan-Keratin, samples underwent secondary tyramide signal amplification. Images were analyzed using Fiji / ImageJ2 (V. 2.3.0) and each sample group was standardized together for brightness and contrast against a secondary-antibody only negative control.

### Contrast enhanced 3D microtomography

We scanned formalin-fixed whole kidneys stained for 2 hours in 4% OsO4 solution (Sigma-Aldrich, St. Louis, MO, USA) using a previously developed method ^31^, at 6 μm isotropic voxel size with high-resolution microtomography (μCT50; Scanco Medical, Brüttisellen, Switzerland) at energy level of 70 keV, and intensity of 57 μA. The tissue volume for each envelope was determined by segmenting all gray-scale images using a fixed Gaussian filter and threshold for all data. Cortical vascular density was determined by inverting the segmented images within the cortex boundaries. Representative images were generated as previously shown ^32,33^ using a heat-map representation of vascular diameter.

### Vascular Density Quantification

Paraffin embedded kidney sections were processed in batch for immunofluorescence staining using the antibodies EMCN and PDPD. Immunofluorescent images were captured on a Ti2 Widefield microscope (Nikon, Tokyo, Japan) at 20x magnification using standardized imaging parameters for each sample group. The images were then processed using the Vessel Analysis plugin for Fiji software (http://imagej.net/Vessel_Analysis). For each preprocessed image, 2-3 regions in the kidney cortex were manually encircled to analyze kidney peritubular capillary and lymphatic density while excluding any glomeruli. The vascular density was measured as a ratio of vasculature area to total area of the encircled kidney cortex surface.

### Statistical analyses

Quantitative data are shown as mean standard error of the mean (SEM). Statistical significance of quantitative results was evaluated using Student’s t-test using GraphPad Prism (version 9.50; www.graphpad.com). *P* values of less than 0.05 were considered statistically significant.

## Acknowledgements

We thank Dr. Guillermo Oliver and Dr. Wanshu Ma for the *VegfcGOF* mouse line. This publication was made possible through core services and support from the Northwestern University George M. O’Brien Kidney Research Core Center (NU GoKidney), an NIH/NIDDK funded program (P30 DK114857). Histology services were provided in part by the Northwestern University Mouse Histology and Phenotyping Laboratory which is supported by NCI P30-CA060553 awarded to the Robert H Lurie Comprehensive Cancer Center. Imaging work was performed at the Northwestern University Center for Advanced Microscopy generously supported by CCSG P30 CA060553 awarded to the Robert H Lurie Comprehensive Cancer Center. Serum BUN measurements were performed by the UAB-USCD O’Brien Center for Acute Kidney Injury Research and are supported by a P30 grant (DK 079337) from the National Institute of Diabetes and Digestive and Kidney Diseases (NIDDK). Dr. Donnan acknowledges his VA employment as a Staff Physician, Medical Service, at Jesse Brown VA Medical Center, Chicago, IL.

## Author Contributions

MD and SQ contributed to the design of the experiments. Animal experiments and histology were performed by MD and DD. Micro-CT imaging was performed by VD. MD, DD, VD, and SQ contributed to analysis of the data. The manuscript was written by MD and SQ. SQ supervised the study. All authors contributed to the review and approval of the manuscript.

